# Regulation of Mechanosensor PIEZO Channel Trafficking in *C. elegans* Germline

**DOI:** 10.64898/2026.08.18.745541

**Authors:** Milagros Rincon Paz, Yooseong Wang, Elva Shao, Sophia Monsalve, Xiaofei Bai

## Abstract

PIEZO proteins are essential mechanotransductive ion channels, yet mechanisms governing their subcellular localization and membrane trafficking remain elusive. Here, we leverage the *C. elegans* germline as an *in vivo* imaging platform to decipher the molecular networks directing PIEZO channel dynamics. High-resolution confocal imaging of endogenously tagged PEZO-1, the *C. elegans* ortholog, reveals that PEZO-1 compartmentalizes with caveolae protein CAV-1 and the recycling endosome regulator RAB-11. Depletion of *rab-11* or the t-SNARE *syx-4* severely impairs the translocation of PEZO-1-positive vesicles to the plasma membrane, establishing a reliance on conserved, RAB-11-dependent machinery. Furthermore, we demonstrated that PEZO-1 structural integrity is essential for vesicle formation and transport; truncating either the N-terminal transmembrane domains or the C-terminal ion pore induces aberrant vesicle morphology and arrests trafficking. Crucially, PEZO-1 vesicle formation and transport are non-autonomously modulated by reproductive signals requiring male sperm or major sperm protein (MSP) signaling. Finally, introducing conserved disease-associated pathogenic PIEZO mutations markedly suppresses its cytosolic and plasma membrane expression. Collectively, our findings define a fundamental cytological framework regulating PIEZO channel dynamics, shedding light on the molecular etiology of PIEZO-associated channelopathies.

## Introduction

PIEZO proteins are essential mechanotransductive ion channels with remarkably diverse physiological roles^1–9^. Structurally, these channels assemble as homotrimers to form a distinct propeller-like or semi-flattened bowl-shaped architecture^10–12^. Their functional output depends on the precise embedding of this trimeric complex within the membrane lipid bilayer. At the target membrane, mechanical stimuli trigger conformational changes in the channel module to open the pore. This allows non-selective cations, including Ca^2+^, to permeate into the cytosol and activate downstream signaling pathways. Despite their physiological prominence, the precise molecular mechanisms directing the trafficking and dynamic sorting of PIEZO channels to their target membranes remain elusive.

When trafficking or surface expression of essential ion channels is disrupted, the resulting cellular dysfunction can lead to severe disorders, collectively known as channelopathies. Under normal physiological conditions, the canonical vesicular trafficking machinery governs membrane protein transport, insertion, membrane fusion, or internalization, serving as a primary determinant of surface channel density^13–15^. For instance, transient receptor potential (TRP) channels rely on rapid, stimulus-responsive vesicle mobilization to modulate their membrane expression^16,17^. Beyond the downstream transport machinery, proper protein folding and oligomeric complex assembly are equally critical for driving nascent protein exit from the endoplasmic reticulum into the cytosol and for their subsequent cytosolic transport and membrane delivery. Consequently, disrupting either the core trafficking machinery or the channel’s structural integrity can reduce functional membrane populations, directly driving pathogenic outcomes. This is well exemplified by the K^+^ channel Kv11.1, in which pathogenic mutations impair both folding and subcellular trafficking, leading to cardiac arrhythmias and long QT syndrome^18^. Deciphering the regulatory networks governing the trafficking of integral membrane proteins is therefore crucial. Uncovering these pathways not only clarifies how channel proteins maintain homeostasis across cytosolic and membrane compartments but also deepens our understanding of the molecular mechanisms underlying these severe channelopathies.

The *C. elegans* germline provides a powerful *in vivo* imaging platform for studying protein trafficking, transport, docking, and membrane fusion, owing to its transparent anatomy and highly conserved cellular machinery^19^. The sole *PIEZO* ortholog, *pezo-1*, encodes 14 distinct splice variants that generate 12 unique proteins. Notably, four of these isoforms lack the hydrophobic N-terminal transmembrane region, also known as the blade domain. Because these N-terminal domains are predicted to drive membrane sorting and targeting, their absence may suppress channel activity by impeding efficient delivery to the cellular membrane^11,12,20–22^. In contrast, the remaining eight elongated isoforms require precise spatiotemporal regulation to achieve functional membrane expression, though the exact cellular pathways governing this process remain undescribed ^21–23^. Furthermore, whether intrinsic structural features or pathogenic mutations affect homomeric assembly and subsequent trafficking is completely unknown. Thus, defining both the accessory proteins involved and the interactions between PEZO-1 monomers is critical to uncovering the foundational mechanisms of PIEZO channel trafficking.

Our previous work revealed that *pezo-1* is indispensable for ovulation and sperm attraction, possibly by coordinating responses to attractive cues, including F-series prostaglandins released by mature oocytes and major sperm protein (MSP) secreted by sperm. Intriguingly, within the *C. elegans* reproductive tract, the membrane secretion and endocytic retrieval of oocyte receptors, such as the Eph receptor VAB-1, are strictly synchronized with sperm availability. This demonstrates a tight functional link between oocyte receptor sorting and sperm-derived signaling inputs, such as the MSP ligands^24^. Therefore, we hypothesize that PEZO-1 trafficking regulation is similarly tied to reproductive signaling and is non-autonomously regulated by inter-tissue signaling exchanges within the reproductive tract.

To examine PIEZO channel transportation and dynamics in the germline, we generated a suite of endogenously tagged PEZO-1 fluorescent reporter lines using CRISPR/Cas9 gene editing. Using these imaging tools combined with high-resolution confocal microscopy, we uncover intricate relationships between PEZO-1 channel trafficking, structural architecture, and extracellular gamete cues. Our data reveal that the channel is dynamically sorted via intracellular membrane trafficking pathways regulated by specific endosomal markers, such as RAB-11, and target membrane fusion SNARE proteins. Furthermore, we demonstrate that overall structural integrity, specifically within the N-terminal transmembrane domains or the C-terminal ion pore, is mandatory for appropriate vesicle morphology and transport, as well as protein expression in the cytosol and membrane. We show that this entire localization pattern is also non-autonomously modulated by external gamete signaling and the soluble major sperm protein (MSP) ligand. Finally, we demonstrate that introducing conserved disease-associated pathogenic mutations dramatically suppresses channel expression across both cytosolic and plasma membrane compartments. In summary, our work established a robust *in vivo* imaging platform to assess PIEZO channel dynamics and local expression. These findings uncover a regulatory network that governs PIEZO channel-mediated cellular homeostasis and provide critical insights into the molecular mechanisms underlying PIEZO-related channelopathies.

## Results

### PEZO-1 localizes to cytosolic vesicles in the germ cells

To characterize the regulatory mechanism governing PIEZO channel membrane trafficking and expression pattern *in vivo*, we previously generated a series of endogenously tagged fluorescent reporters at the endogenous *pezo-1* locus using CRISPR/Cas9, which allowed accurate visualization of PEZO-1 expression and intracellular trafficking in living animals. Leveraging the transparent anatomy and highly conserved cellular machinery of *C. elegans*, we first characterized the intracellular localization and trafficking of PEZO-1 in the germline. N-terminal tagged GFP::PEZO-1 (Fig.1A), and C-terminal tagged PEZO-1::mScarlet (Fig.1B) labeled intracellular vesicles spread throughout the adult hermaphrodite germline (Fig.1A-C). The diameters of these PEZO-1-positive vesicles ranged from approximately 0.5 µm to 1.5 µm (Fig.1G), with a subset displaying distinct, ring-like structures (yellow and red insets, Fig.1A-B). Live imaging of PEZO-1 trafficking revealed that these positive vesicles were transported from the perinuclear region to the target plasma membrane, where GFP::PEZO-1 was tethered and fused with the membrane (yellow arrows, Fig.1C). These data demonstrate that PEZO-1 can efficiently traffic to the plasma membrane from the sites of synthesis within germline cells.

**Figure 1.**
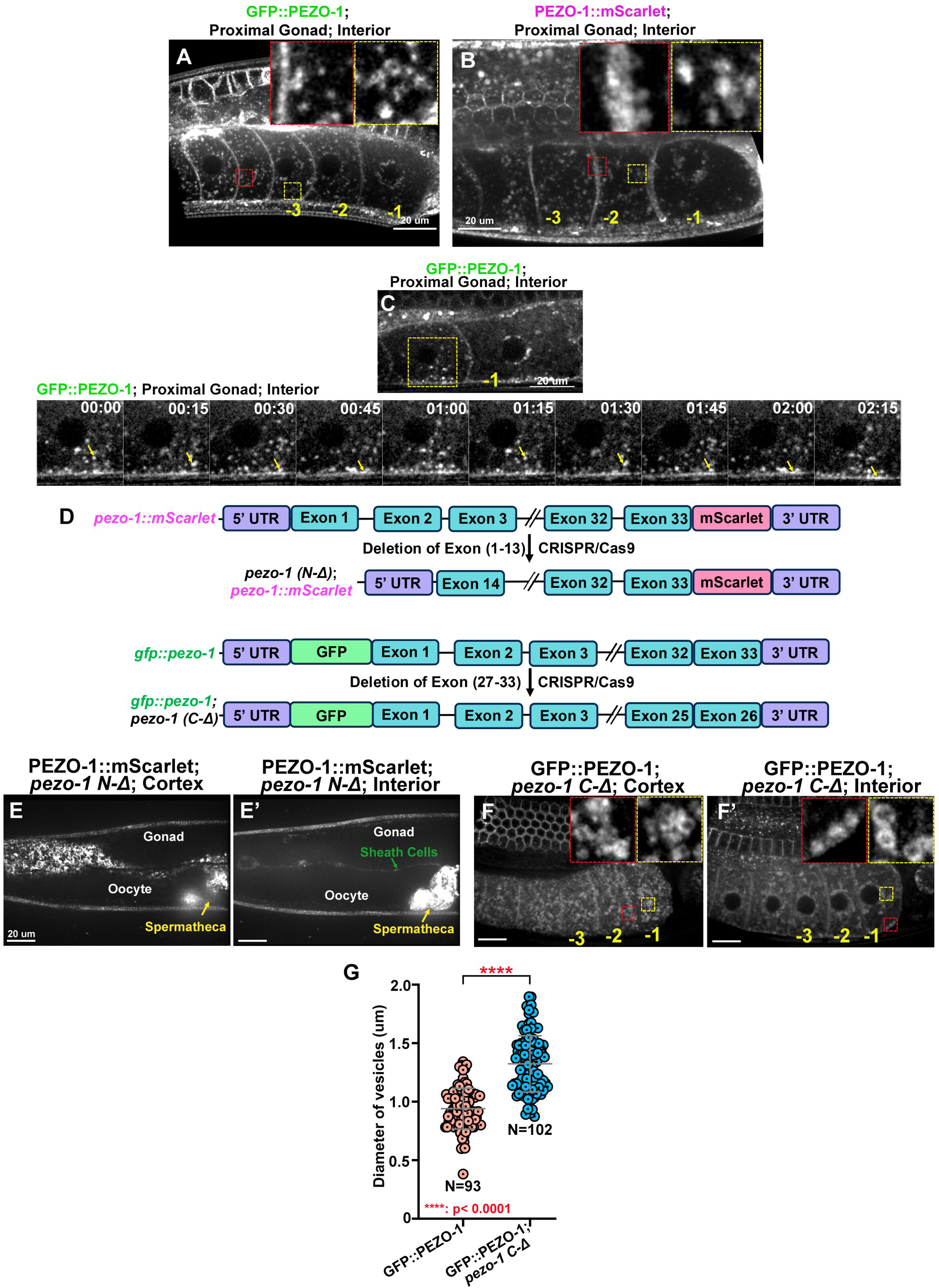
PEZO-1 localizes to cytosolic vesicles in the germ cells and requires structural integrity for proper trafficking and localization. (A-C) Representative images of endogenously tagged GFP::PEZO-1 (A, C) and PEZO-1::mScarlet (B) in the living adult hermaphrodite germline. Images were acquired in the proximal gonad (A-C) and visualized at the central focal planes. Inserts show magnified views of the selected region highlighted by yellow and red boxes. (C) Representative images and kymograph (F) of GFP::PEZO-1 positive vesicle trafficking in the oocytes. The timing of each step is labeled in the top-right corner in minutes and seconds. (D) Schematic of the two deletion alleles were generated in two PEZO-1 fluorescent reporter strains. (E-E’) The fluorescent signals of PEZO-1::mScarlet were only observed in somatic tissues but not in the germline in *pezo-1 N-Δ.* (F-F’) In contrast, C-terminal truncated GFP::PEZO-1 was observed throughout the germline. The truncated GFP::PEZO-1-containing vesicles are abnormally enriched at the cellular cortex and peri-nuclear region. Inserts show magnified views of the aggregated vesicles (yellow and red inserts). (G) Quantification of both integral GFP::PEZO-1 and truncated GFP::PEZO-1 vesicles diameters in the germline. Scale bars are indicated in each panel. P-values: **** <0.0001 (t-test) (G).

The *pezo-1* gene encodes 14 splice variants that generate 12 unique proteins. Four of these isoforms lack the hydrophobic N-terminal tandem transmembrane domains known as “blade” domains^11,12^. Because these N-terminal domains are predicted to anchor the PEZO-1 trimeric complex into the lipid bilayer, their absence may alter channel density at the plasma membrane^11,12,21,22,25^. To determine whether the structural integrity of PEZO-1 is required for proper channel trafficking and regulation, we generated two truncated alleles in our fluorescent reporter backgrounds using CRISPR/Cas9 editing. The first allele was engineered into the PEZO-1::mScarlet background with a deletion of exons 1-13 and flanking introns, and was named as *pezo-1 N-Δ* (Fig.1D). The second allele consists of a deletion of the final seven exons (27-33) and intervening introns, named as *pezo-1 C-Δ*, in the GFP::PEZO-1 strain (Fig.1D).

In *pezo-1 N-Δ* mutants, the truncated PEZO-1::mScarlet protein was widely expressed in somatic tissues, including the somatic sheath cells and the spermatheca (arrows in Fig.1E-E’). However, no fluorescent signal was detected within the oocytes or the proximate germline (Fig.1E-E’), suggesting that the N-terminal transmembrane segments are indispensable for PEZO-1 expression or stability in the germline. Conversely, deletion of the C-terminal ion pore module domains (*pezo-1 C-Δ*) causes aberrant vesicle morphology and disrupted trafficking (Fig.1F-F’). The diameters of these C-terminally truncated GFP::PEZO-1 vesicles were significantly enlarged compared to those containing full-length GFP::PEZO-1 (Fig.1G). Furthermore, these truncated vesicles aggregated prominently at the cortical membrane (yellow inserts, Fig.1F) as well as in the perinuclear region (red and yellow inserts, Fig.1F’), suggesting that the C-terminal domains are critical for maintaining proper vesicle morphology and directing intracellular trafficking. Collectively, these findings suggest that the delivery of PEZO-1 to the target membranes within the germline strictly requires the structural integrity of both its N- and C-terminal domains.

Finally, we tested whether germline PEZO-1 is synthesized autonomously within the germline, or instead originates from the surrounding somatic gonad and is endocytosed by somatic sheath cells. We leveraged the auxin-inducible degradation (AID) system to selectively degrade PEZO-1 within either the entire germline or the soma (Supplemental Fig.1A)^11^. Upon auxin exposure, GFP::PEZO-1::AID fluorescent intensities within the germline and on the oocyte plasma membrane were significantly reduced in the germline-specific AID strain (Supplemental Fig.1B-C). A concomitant and significant reduction of positive vesicles in the cytosol or cell cortex was also observed under the auxin treatment (Supplemental Fig.1C-D). In contrast, GFP::PEZO-1::AID fluorescent intensities or vesicle patterns were not visibly altered in the soma-specific AID strain, which thoroughly degrades the somatic GFP::PEZO-1::AID pool, including within the spermatheca (yellow arrow in Supplemental Fig.1E-F). These data demonstrate that the PEZO-1 vesicle population is synthesized endogenously within germline cells rather than being imported from surrounding somatic tissues.

### PEZO-1 trafficking is regulated by endosomal trafficking regulators in the germline

To gain insight into the molecular mechanisms governing PEZO-1 trafficking, we investigated whether PEZO-1-positive vesicles associate with specific components of the intracellular membrane trafficking machinery. We first focused on RAB-11, a small GTPase that plays an essential role in regulating recycling endosome trafficking and vesicle fusion at the target membrane ^26,27^. In animals co-expressing mScarlet::PEZO-1 and RAB-11::GFP, extensive colocalization of the two fluorescent reporters was observed on both cortical and interior vesicles throughout the germline (Fig.2A-B). In contrast, we did not detect any significant colocalization between GFP::PEZO-1 and the early endosomal marker mCherry::RAB-5 (Supplemental Fig.1A-B). This divergence indicates that the steady-state intracellular pool of PEZO-1 does not dynamically reside in early endocytic pathways but is instead selectively enriched in the RAB-11-associated recycling endosome compartment.

**Figure 2.**
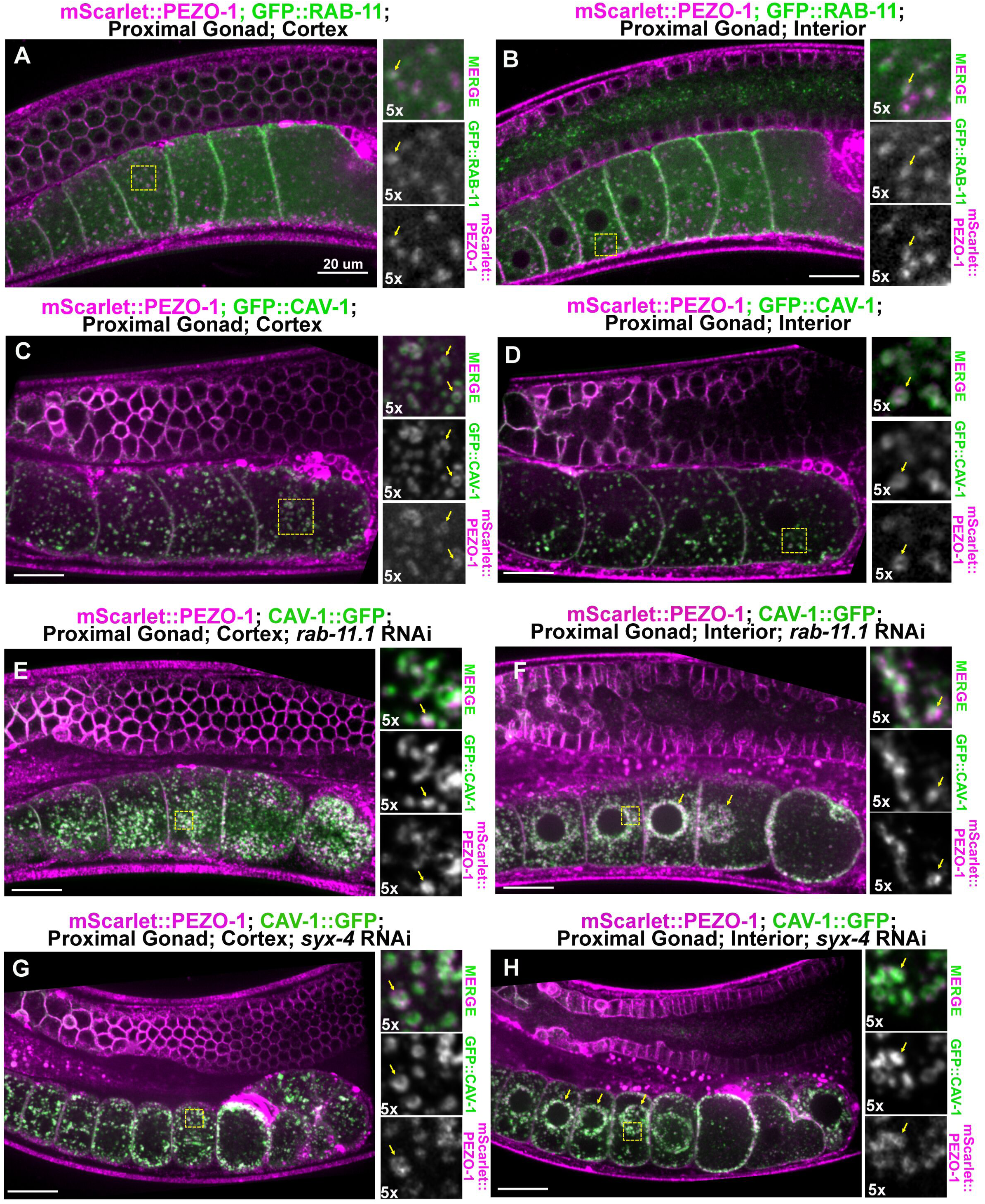
PEZO-1 is enriched within the specialized RAB-11 and CAV-1 positive recycling endosomal compartment. (A-B) mScarlet::PEZO-1 (magenta) colocalized with the recycling endosome marker GFP::RAB-11 (green) positive compartment. The right panels show magnified views of the area marked by yellow squares. Images were also acquired in both the cell cortex (A) and the interior cytosol (B) of proximal oocytes, visualized at the central focal planes. (C-D) Representative images showing spatial colocalization between mScarlet::PEZO-1 (magenta) and the caveolar vesicle marker GFP::CAV-1 (green) on characteristic ring-shaped cytosolic vesicles. Yellow arrows in the right insets indicate precise vesicular colocalization at the cortical (C) and interior (D) regions of proximal oocytes. (E-H) The localization pattern of mScarlet::PEZO-1 (magenta) and CAV-1::GFP (green) under different RNAi conditions. Depletion of both *rab-11.1* (E-F) and *syx-4* (G-H) by RNAi, leading to abnormal accumulation of cargo vesicles underneath the plasma membrane at the cell cortex (yellow insert, E, G) and peri-nuclear area (yellow arrows and inserts, F, H). Scale bars are indicated in each panel.

Because recycling endosomes often serve as sorting stations that direct diverse transmembrane cargos to the same target membrane sites, we next examined whether PEZO-1 shares transport carriers with other known membrane-associated proteins. In mammalian systems, the structural protein, Caveolin-1 (CAV1), which forms flask-shaped caveolar membrane invaginations, colocalizes with PIEZO1 to modulate its Ca^2+^ dependent mechanical activation^28^. In *C. elegans*, the caveolin-1 ortholog CAV-1 is synthesized in the germline, generated by the Golgi apparatus, and transits through the endomembrane system to fuse with the plasma membrane in maturing oocytes^29^. We therefore imaged mScarlet::PEZO-1 alongside CAV-1::GFP to determine if they share a common transport pathway. Similar to the pattern observed with RAB-11::GFP, CAV-1::GFP colocalized extensively with mScarlet::PEZO-1 on characteristic ring-shaped vesicles in the germline, particularly within the proximal oocytes (Fig.2C-D). Additionally, we observed a parallel vesicular colocalization pattern with the Eph receptor VAB-1::GFP, which also undergoes endocytic recycling in the germline (Supplemental Fig.2C-D). These spatial overlaps demonstrate that PEZO-1 resides with CAV-1 and VAB-1, trafficking from the same RAB-11-positive recycling endocytic compartment within the *C. elegans* germline.

To determine if the transport of these PEZO-1 carrier vesicles to the plasma membrane directly depends on RAB-11 activity, we performed germline-specific RNAi against *rab-11.1* in animals co-expressing mScarlet::PEZO-1 and CAV-1::GFP. Knockdown of *rab-11.1* significantly disrupted the distribution of these vesicles, causing them to stall and abnormally cluster both near the cell cortex (yellow inserts in Fig.2E) and within the peri-nuclear region (yellow arrows in Fig.2F). Notably, despite this severe transport arrest, CAV-1::GFP and mScarlet::PEZO-1 remained closely associated on the membrane of these stalled, ring-shaped vesicles (right channels in Fig.2E-F). This finding demonstrates that RAB-11 is indispensable for the long-range transport of PEZO-1-positive compartments to the plasma membrane, but is not required for the initial sorting of PEZO-1 and CAV-1 into the same vesicular carriers.

Finally, we examined the terminal step of this trafficking itinerary by analyzing the physical docking and fusing of PEZO-1 vesicles at the target membrane sites. This terminal tether-docking step typically requires the target-membrane-localized SNARE (t-SNARE) proteins to mediate vesicle docking and bilayer fusion^27,30^. We hypothesized that if PEZO-1 vesicles utilize a conventional exocytosis itinerary to integrate into the plasma membrane after their RAB-11 mediated transit, depletion of the t-SNARE *syx-4* (syntaxin-4) would block their terminal fusion. Consistent with this hypothesis, *syx-4*(RNAi) disrupted vesicle integration at the cell cortex (Fig.2G-H). In the *syx-4* RNAi-treated animals, mScarlet::PEZO-1 vesicles failed to fuse with the plasma membrane and instead accumulated in the sub-cortical cytoplasm beneath the target membrane (Fig.2G-H, yellow arrows in Fig.2H).

Taken together, these data show that PEZO-1 resides in a specialized recycling endosomal compartment marked by RAB-11 and CAV-1. Forward transit of these carrier vesicles through the cytoplasm requires RAB-11, while their final integration and fusion with the target plasma membrane relies on the canonical SNARE machinery.

### The PEZO-1 trafficking is regulated by the presence of sperm

Our previous work demonstrated that PEZO-1 functions as a positive regulator of oocyte ovulation and sperm attraction back to the spermatheca^25,31,32^. Dysfunction of PEZO-1 causes a severe reduction in both self-sperm count and ovulation rate^25,32^. Having established that PEZO-1 utilized the aforementioned endosomal pathway to reach the oocyte membrane, we next investigate whether physiological cues within the reproductive tract, particularly sperm availability, modulate this trafficking pathway. To test this, we crossed the GFP::PEZO-1 reporter into a temperature-sensitive *fem-1(hc17)* mutant background, which completely lacks spermatogenesis and develops exclusively as females when maintained at the non-permissive temperature of 25°C^33^,

In unmated *fem-1(hc17)* females at 25°C, where sperm are absent from the reproductive tract, GFP::PEZO-1 exhibited a progressive and profound cortical enrichment in proximal oocytes (Fig.3A-B). GFP::PEZO-1 became highly trapped within cortically localized vesicles (yellow arrows and yellow inserts in Fig.3A-B), and was predominantly restricted beneath or at the plasma membrane in the steady state (red inserts in Fig.3A-B). In contrast, when *fem-1(hc17)* females were mated with wild-type males, the introduction of sperm fully restored the GFP::PEZO-1 localization and vesicle patterns identical to those observed in wild-type hermaphrodites (Fig.3C-E). This striking reversal indicates that sperm availability serves as a critical physiological cue for maintaining proper, dynamic PEZO-1 vesicle trafficking and turnover in the germline.

**Figure 3.**
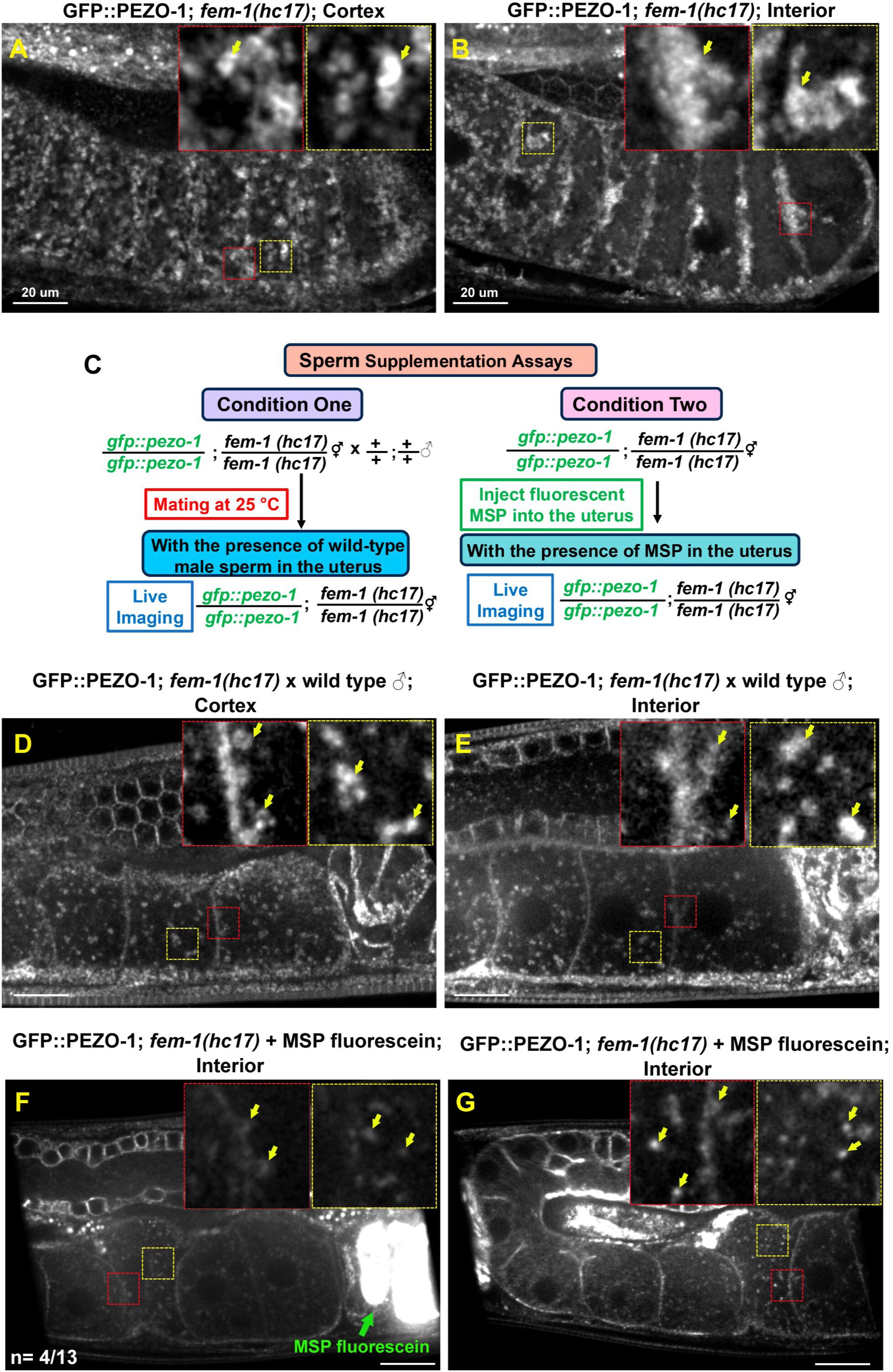
Dynamic PEZO-1 vesicular trafficking requires the sperm-derived signaling. (A-B) GFP::PEZO-1 in unmated sperm-depleted *fem-1(hc17)* female animals at non-permissive temperature (25 °C). In the absence of sperm, GFP::PEZO-1 accumulated in static, trapped vesicles at the cellular cortex (red and yellow inserts, A) or directly adjacent to the plasma membrane (red and yellow inserts, B). Top-right insets show magnified views of the selected regions, with GFP::PEZO-1 enrichment in the membrane (red squares) or the cortex (yellow squares). (C) Experimental workflow schematic illustrating the male mating and purified Major Sperm Protein (MSP) microinjection supplementation assays. (D-E) Representative images of *fem-1(hc17)* females post-mating with wild-type males. The introduction of mature sperm restores normal cytosolic vesicle distribution and alleviates abnormal cortical trapping. The top right insets show magnified views of the cortex (yellow squares) and plasma membrane (red squares). (F-G) Injection of purified fluorescein-tagged MSP in the uteri of the unmated *fem-1(hc17)* females with GFP::PEZO-1. The fluorescein-tagged MSP ligand successfully translocates through the uterus to enrich at the spermatheca (green arrow in F). Supplemental MSP signaling is sufficient to partially clear the aberrant cortical trapping of GFP::PEZO-1, restoring dynamic cytosolic trafficking patterns in a subset of treated animals (red and yellow squares). Scale bars are indicated in each panel.

In *C. elegans*, signaling from the sperm to the oocyte is primarily mediated by the Major Sperm Protein (MSP), hormone-like signaling ligands secreted by proximal spermatozoa^34–36^. To determine if MSP signaling is sufficient to modulate PEZO-1 trafficking in the absence of whole sperm, we microinjected purified, fluorescein-labeled MSP into unmated *fem-1(hc17)* females^37^. Remarkably, approximately 30% (n= 4/13) of the MSP-injected animals displayed a distinct elevation of the abnormal GFP::PEZO-1 accumulation at the plasma membrane and cell cortex, partially restoring its dynamic cytosolic trafficking pattern (Fig. 3C, F-G). Collectively, these data demonstrate that sperm-derived MSP signaling actively remodels PEZO-1 intracellular trafficking, highlighting a potential feedback mechanism by which PEZO-1 senses sperm availability to coordinately promote oocyte ovulation.

### PEZO-1 localizes to the cortical granules and undergoes exocytosis in anaphase I embryos

To determine whether the intracellular transport mechanisms identified in the adult germline represent a conserved cellular strategy for the PEZO-1 channel, we extended our analysis to another specialized membrane trafficking event, cortical granule exocytosis (CGE). Cortical granules (CGs) are large secretory vesicles, approximately 1 µm in diameter, that rely on canonical trafficking regulators, including RAB-11, for their transport and exocytosis in *C. elegans* meiotic embryos ^38,39^. Crucially, cortical granule exocytosis is a highly synchronized, rapid, and developmentally regulated vesicle-fusion event that occurs immediately following meiotic anaphase I, providing an ideal physiological window to capture real-time channel delivery kinetics.

Intriguingly, we found that mScarlet::PEZO-1 co-localized extensively with two established cortical granule markers, RAB-11::GFP and CAV-1::GFP^29,40^, on characteristic ring-shaped cortical granules within meiotic I embryos (Fig.4A-B, yellow arrows and insets, right panels). To capture the real-time dynamics of this compartment, we utilized high-temporal resolution live imaging to track individual vesicles labeled with either GFP::PEZO-1 (Fig.4C-G), or mScarlet::PEZO-1 (Supplemental Fig.3A-D). These assays demonstrated that PEZO-1-positive compartments undergo stereotypic exocytosis during anaphase I. Specifically, real-time tracking of a single GFP::PEZO-1-positive granule captured the precise moment of vesicle integration and bilayer fusion with the target plasma membrane. Following cargo release, the GFP::PEZO-1 signal smoothly merged with and spread into the plasma membrane (Fig. 4G), demonstrating that these PEZO-1-positive compartments belong to the maternal population of cortical granules destined for meiotic exocytosis.

**Figure 4.**
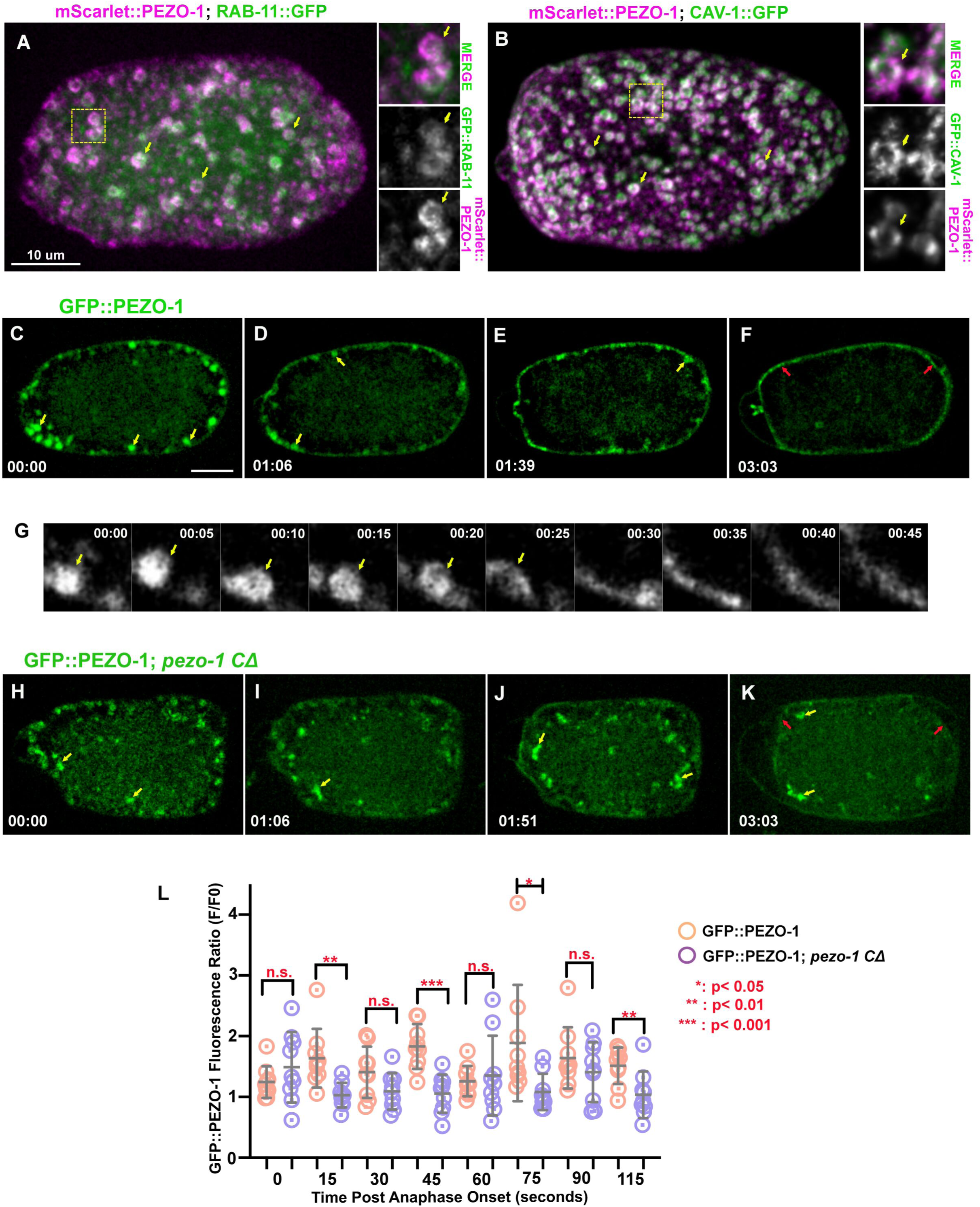
PEZO-1 localizes to maternal cortical granules and undergoes exocytosis during embryonic anaphase I. (A) Surface plane image of a meiotic I embryo coexpressing mScarlet::PEZO-1 (magenta) and RAB-11::GFP (green), showing colocalization on the cortical granules. The yellow arrow in the right panels points to a representative double positive vesicle of mScarlet::PEZO-1 (magenta) and RAB-11::GFP (green). (B) A similar colocalization of cortical granules was observed in animals expressing mScarlet::PEZO-1 (magenta) and CAV-1::GFP (green). The yellow arrow in the right panels denotes a single vesicle of mScarlet::PEZO-1 (magenta) and RAB-11::GFP (green) in the cytosol. The right panels show magnified views of the selected region (yellow squares). (C-F) High-temporal-resolution live imaging captures the real-time dynamics of endogenous GFP::PEZO-1 (green) during cortical granule exocytosis. (G) Representative kymographic images show a single vesicle tether, fusion, and exocytosis during anaphase I. GFP::PEZO-1 was associated with the plasma membrane after exocytosis. (H-K) Time-lapse imaging of meiotic embryos expressing C-terminally truncated *pezo-1(C-Δ)*. The truncated channel leads to severe structural defects, resulting in static, highly aggregated vesicle clusters in the cytosol and reduced plasma membrane expression of GFP::PEZO-1 (yellow arrows). (L) Quantification of the post-exocytic plasma membrane fluorescence intensity over time, showing a significant reduction in surface channel population in the C-terminal truncated embryos after the onset of anaphase I. Scale bars are indicated in each panel. P-values: *: p <0.05 (t-test); **: p <0.01 (t-test); ***: p <0.001 (t-test).

We next resolved the post-fusion dynamics of the PEZO-1 channel relative to its trafficking machinery. Following vesicle fusion, both GFP::PEZO-1 (red arrows in Fig.4F) and mScarlet::PEZO-1 (red arrows in Supplemental Fig.3D) remained stably associated with the plasma membrane. To directly evaluate the coordination between this membrane fusion event and the behavior of the upstream transport machinery, we filmed meiotic embryos coexpressing mScarlet::PEZO-1 and RAB-11::GFP. Prior to exocytosis, RAB-11::GFP fully co-localized with mScarlet::PEZO-1 positive granules (Supplemental Fig.3A-C). Upon exocytosis, however, the RAB-11::GFP signal rapidly dissociated from the cortex and diminished in the cytosol, whereas mScarlet::PEZO-1 remained persistently anchored at the target plasma membrane (red arrows in Supplemental Fig.3D). This distinct kinetic divergence supports a model where RAB-11 acts transiently as a delivery factor, promoting the translocation and stable incorporation of PEZO-1 into the newly remodeled embryonic plasma membrane.

Finally, to verify whether the structural requirements for embryonic PEZO-1 trafficking recapitulate our observations in the adult germline, we evaluated meiotic embryos expressing the truncated PEZO-1 reporters. Paralleling our germline findings, the C-terminally truncated GFP::PEZO-1 caused severe vesicular morphology and transport defects, frequently forming aberrant, static vesicle clusters that aggregated within the cytosol (yellow arrows in Fig. 4H-K). This transport failure resulted in a significant reduction in the plasma membrane-bound GFP::PEZO-1 fluorescence following the exocytic wave (red arrows in Fig. 4K and Fig. 4L). In contrast, deletion of the N-terminal domains completely abolished detectable vesicular fluorescence within the meiotic embryos, precluding further evaluation of how these N-terminal domains affect the exocytosis step. Collectively, these data demonstrate that the efficient delivery, fusion, and steady-state expression of PEZO-1 at the embryonic plasma membrane depend on channel structural integrity and on the conserved cortical granule exocytosis pathway.

### Pathogenic PEZO-1 mutations alter intracellular localization and protein expression in *C. elegans*

Previous clinical studies have demonstrated that numerous pathogenic mutations in ion channels and other transmembrane proteins, including in Kv11.1 (the voltage-gated potassium channel associated with long QT syndrome) and NPHS1 (associated with congenital nephrotic syndrome), primarily cause disease by disrupting protein folding, trafficking, or membrane expression^41–43^. Here, we sought to test whether conserved, disease-associated mutations within the C-terminal modules of PIEZO channels alter channel trafficking or membrane expression *in vivo*. By leveraging our established imaging system for PEZO-1 transport, we aimed to shed light on the molecular mechanisms underlying PIEZO-associated channelopathies. We introduced a suite of conserved pathogenic mutations into the C-terminus of our endogenous GFP::PEZO-1 reporter strain using CRISPR/Cas9 genome editing. These variants correspond to human PIEZO1 substitutions (p.R2456H and p.R2488Q) implicated in Dehydrated Hereditary Stomatocytosis (DHS) and Generalized Lymphatic Dysplasia (GLD), and PIEZO2 substitutions (p.R2686C/H and pR2718L/P) causing Distal Arthrogryposis Type 3 (DA3) and Type 5 (DA5). Additionally, we generated compound double-mutant strains by combining *pezo-1*[R2373C/H] and *pezo-1*[R2405L/Q] to determine whether combining these structural perturbations would exacerbate trafficking defects. Notably, these mutations cause severe physiological defects in *C. elegans*, including a significant reduction in brood size, suggesting that the worm mutations may have similar pathological effects on ion channel activity^44^.

Upon isolating homozygous mutants, we observed that most strains exhibited a profound reduction in total PEZO-1 expression across both cytosol and plasma membrane within the germline (Fig. 5A-J). For specific variants, such as *pezo-1*[R2405P] and *pezo-1*[R2405L], PEZO-1-positive vesicles were barely detectable (Fig.5A, D-E, H). This severe loss of fluorescence was recapitulated in somatic tissues, including the pharyngeal-intestinal valve and the spermathecal cells, where wild-type PEZO-1 acts as a functional mechanosensitive ion channel to support essential physiological processes, including pharyngeal pumping and ovulation (Supplemental Fig.4A-H, Supplemental Fig.5A-H) ^21,25,31,44,45^. Collectively, our observations suggest that these missense mutations within the C-terminal module autonomously destabilize the PEZO-1 complex, leading to aberrant trafficking or accelerated degradation before the channel reaches its target compartment.

**Figure 5.**
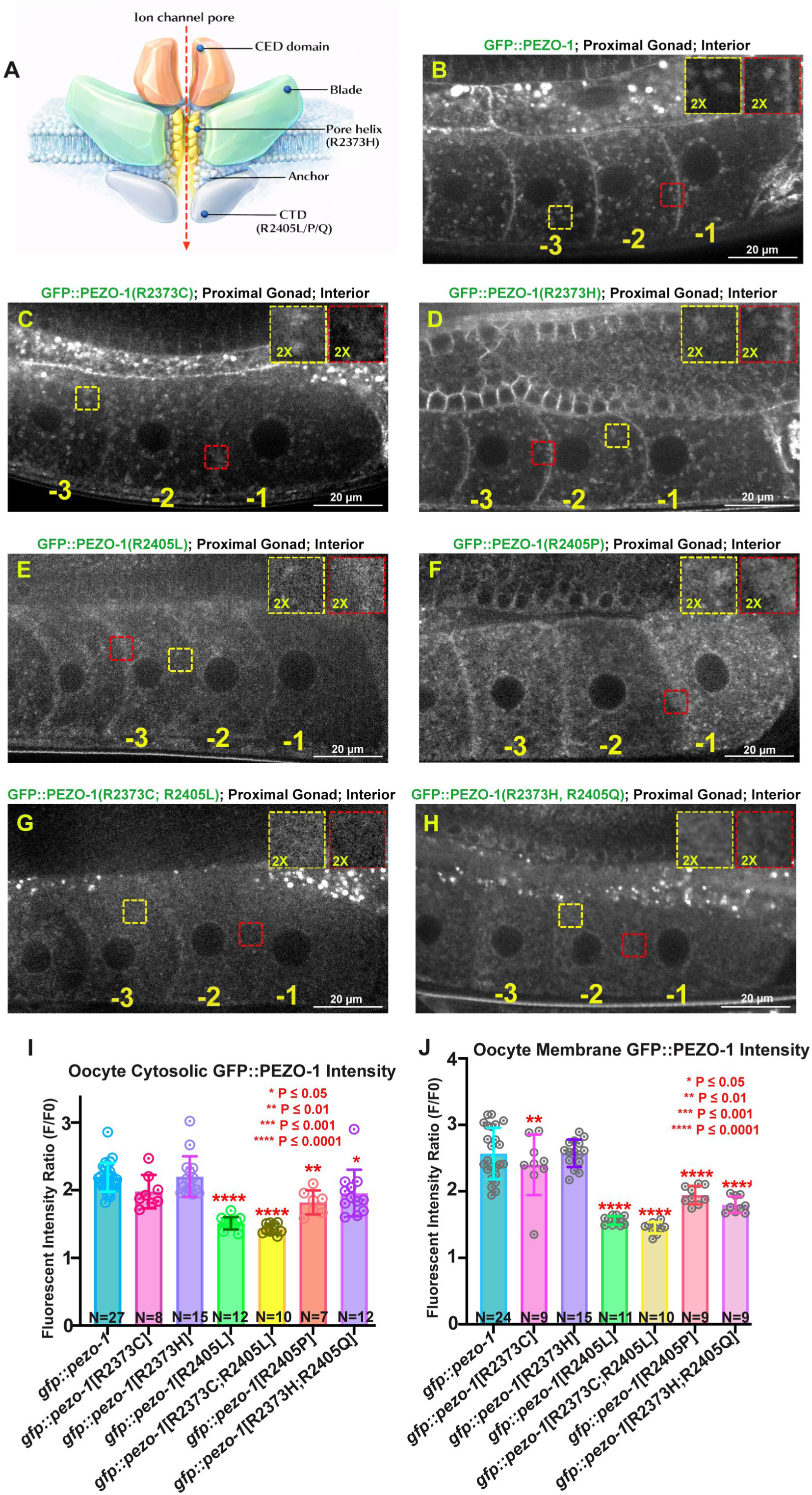
Pathogenic mutations impair PEZO-1 protein stability and membrane localization. (A) Structural schematic diagram of the PEZO-1 mechanosensitive ion channel complex, including the C-terminal extracellular domain (CED), the blade, the anchor, the pore helix harboring the human disease-associated variant site (R2373H/C), and the C-terminal domain (CTD) containing variant sites (R2405L/P/Q). The dashed red arrow marks the ion channel pore axis. (B–H) Representative images show the proximal gonad and oocytes (labeled -3, -2, -1 relative to the spermatheca) in living adult hermaphrodites. (B) The endogenously tagged wild-type GFP::PEZO-1 shows the characteristic ring structure of GFP::PEZO-1-positive vesicles in the oocyte. (C-H) The representative images of each human disease-associated variant in the GFP::PEZO-1 background. The pore helix single mutants R2373C (C) and R2373H (D); the CTD single mutants R2405L (E) and R2405P (F); and the double-mutant combinations R2373C; R2405L (G) and R2373H; R2405Q (H). Yellow and red boxes outline selected regions magnified two times in the upper right insets to highlight differences in intracellular vesicle morphology and membrane enrichment. (I-J) Quantitative analysis of the relative oocyte cytosolic and plasma membrane GFP::PEZO-1 fluorescence intensity ratio across wild-type and indicated single- and double-mutant strains, suggesting that missense variants within the CTD structural core cause a profound loss of both cytosolic vesicle signal and steady-state plasma membrane integration. P-values: *: p <0.05; **: p <0.01; ***: p <0.001; ****: p <0.0001 (One-Way ANOVA with Dunnett’s post-hoc test). Scale bars are indicated in each panel.

To investigate whether these expression and trafficking defects could be bypassed or alleviated by genetic pathways that restore physiological functions, we evaluated the previously identified genetic modifiers in the *pezo-1*[R2405P] mutant. This loss-of-function genetic modifier resides within the WAVE complex gene *gex-3* and suppresses the reduced brood size of the *pezo-1*[R2405P] mutant by modulating actin polymerization defects in the spermathecal cells. We tested whether knocking down *gex-3* by RNAi could alleviate the underlying membrane trafficking or expression defects observed in the *pezo-1*[R2405P] background. However, the *gex-3*(RNAi) treatment failed to restore GFP::PEZO-1 fluorescence or alter its defective distribution (Supplemental Fig.6A-F). This divergence demonstrates that genetic rescue does not occur by repairing upstream subcellular trafficking and expression defects, but rather by potentially optimizing tissue-level function or amplifying the structural signaling efficiency of the scarce residual channel pool at the membrane.

Collectively, these data suggest that pathogenic mutations in the C-terminal module of PEZO-1 fundamentally destabilize channel transport and steady-state expression. Because the C-terminal module contains the pore-forming domain and the trimerization assembly required for proper channel architecture, these mutations likely trigger cellular protein quality-control pathways that compromise protein stability and membrane trafficking *in vivo*. Further studies will determine whether these expression defects originate from a loss of ion-channel-dependent signaling mechanisms, such as localized Ca^2+^ homeostasis, an inability to physically interact with the core membrane-delivery machinery, or severe structural-biophysical defects that disrupt proper PEZO-1 homotrimer assembly.

## Discussion

In this study, we established an *in vivo* imaging system to elucidate the mechanisms that regulate the trafficking and membrane localization of the mechanotransductive ion channel PEZO-1 in the *C. elegans* germline. We show that PEZO-1 membrane delivery depends on a conserved endosomal recycling pathway mediated by the small GTPase RAB-11, which directs the channel from intracellular trafficking compartments to the plasma membrane. During transport, PEZO-1 is co-packaged into RAB-11-positive vesicles together with the caveolar protein CAV-1 and the Ephrin receptor tyrosine kinase VAB-1, suggesting coordinated trafficking of multiple membrane-associated signaling proteins. The final step of membrane delivery requires the SNARE complex component SYX-4, indicating that SNARE-dependent vesicle tethering and fusion are essential for PEZO-1 surface localization. We further demonstrate that efficient trafficking depends on the channel’s structural integrity. Truncation of either the N- or C-terminal regions, as well as patient-derived missense variants within the conserved C-terminal domains, can cause intracellular retention, protein aggregation, or degradation, thereby reducing membrane localization. Finally, we reveal that PEZO-1 surface expression is dynamically regulated by extracellular and inter-tissue signaling cues, demonstrating that physiological signals can actively modulate intracellular trafficking of the PEZO-1 channel.

Although the biophysical properties of PIEZO channels have been extensively investigated, the cellular mechanisms governing their intracellular trafficking, vesicle sorting, and plasma membrane delivery have remained elusive. Our findings identify a trafficking pathway in which RAB-11-dependent recycling endosomes and SNARE machinery cooperate to deliver PEZO-1 to the plasma membrane. Given the exceptionally large size and trimeric architecture of PEZO-1 channels, regulated trafficking is essential to ensure their efficient transport and correct membrane localization. Moreover, the colocalization of PEZO-1 with CAV-1 and VAB-1 suggests that mechanosensory and signaling proteins may be assembled into common transport vesicles before reaching the cell membrane, potentially coordinating the formation of functional macromolecular domains at the plasma membrane.

Our findings align with emerging evidence that Rab GTPase-mediated membrane trafficking serves as an evolutionarily conserved mechanism governing PIEZO channel trafficking and localization. For example, during mammalian cytokinetic abscission, PIEZO1 localizes to the intercellular bridge, where it functions alongside RAB-11-FIP3-positive recycling endosomes to generate local calcium transients that promote vesicle delivery and membrane remodeling prior to daughter cell separation ^46^. Likewise, in *Drosophila* sensory neurons, Piezo channel localization is regulated by another small GTPase, Rab10, which acts upstream of integrin signaling to control post-injury axonal regeneration ^47^.

Our structure-function studies further demonstrate that the intracellular trafficking and stability of PEZO-1 are tightly linked to its structural integrity. Deletion of the hydrophobic N-terminal blade domains, which are crucial for forming the characteristic trimeric complex, resulted in a dramatic reduction in protein expression. This aligns with a recent study showing that disruption of the N-terminal propeller blade domain of PIEZO1 leads to protein misfolding, resulting in channel fragments being retained within the endoplasmic reticulum (ER) ^63^.

Intriguingly, truncating the C-terminal module of PEZO-1 led to abnormal intracellular aggregation rather than rapid proteolytic clearance; this deletion allele produced enlarged, static cytosolic vesicle clusters. Because wild-type PEZO-1 channels possess a curved, propeller-like physical architecture, their active insertion into lipid bilayers exerts a significant mechanical force that can induce membrane curvature^11,12^. We propose that the C-terminal pore domain of PEZO-1 may act as an intrinsic regulator of vesicle homeostasis. Truncating this module likely disrupts normal membrane bending during vesicle budding or maturation, leading to the abnormal fusion of stalled cargo carriers within the cytoplasm. Thus, the PEZO-1 channels may not only act as a passive cargo within recycling endosomes but also actively determine the morphology of the transport vesicles that carry them.

Similarly, missense mutations residing within the conserved C-terminal ion channel module disrupted membrane trafficking and expression, consistent with previous mammalian and clinical studies. This directly recapitulates foundational work demonstrating that PIEZO surface expression is strictly dependent on proper structural assembly and channel functions ^48,49^. Introducing patient-derived missense mutations into either the conserved C-terminal pore module or the stabilizing beam domains compromised protein stability and blocked membrane trafficking ^48–51^. Several of these clinical variants—such as those associated with General Lymphatic Dysplasia (GLD)—exhibited reduced membrane trafficking, triggered recognition by the ER retention machinery, and led to rapid clearance via Endoplasmic Reticulum-Associated Degradation (ERAD)^48^. Together, these findings suggest that structural integrity and proper folding of the PEZO-1 channel are prerequisites for efficient trafficking through the secretory pathway.

In mammalian models, PIEZO channels do not function in isolation; their channel gating, surface density, and mechanical sensitivity are extensively modulated by extracellular chemical ligands and inter-tissue signaling pathways, such as Gi-coupled ligands, polyunsaturated fatty acids, ceramides, and TNF-related apoptosis-inducing ligand ^52–55^. Consistent with this concept, we found that basal PEZO-1 transport and cellular stability in *C. elegans* are dynamically regulated by extracellular cues and inter-tissue signaling, specifically by the presence of sperm or sperm-derived signaling cues. The profound arrest of vesicle trafficking observed in the absence of sperm signaling—and its rapid reversal by externally supplied MSP protein—establishes that external sperm cues actively regulate functional channel retention and transport. This mechanism ensures that these essential mechanosensitive channels are spatiotemporally regulated and are delivered to the plasma membrane only when the local physiological environment demands mechanical coordination, such as fertilization and ovulation, thereby preventing premature or aberrant channel activation.

Collectively, this study provides an extensive *in vivo* dissection of an endogenously regulated trafficking itinerary of the PIEZO channel. While current research remains heavily focused on the electrophysiological kinetics of the PIEZO channels in isolated, cultured cell systems, our discovery of these complex endomembrane pathways, structural integrity checkpoints, and physiological cues offers a much-needed complementary perspective. By uncovering the logistical mechanisms governing the assembly, transport, and spatial distribution of this essential mechanosensor *in vivo*, our work provides additional cellular insights into the pathological etiology of PIEZO-associated channelopathies, particularly those that drive disease through affecting channel trafficking and protein instability rather than altering channel gating and kinetics.

## Materials and Methods

### *C. elegans* strains used in this study

We maintain *C. elegans* on OP50-seeded MYOB plates. AG551 and AG552 were generated by crossing AG404 with WH347 and RT688 hermaphrodites containing RAB-11::GFP and CAV-1::GFP, respectively. F3 adults were screened for the presence of the mScarlet with GFP reporters. Detailed strain information is listed in the supplemental information.

### RNAi treatment

The RNAi feeding constructs were obtained from the Vidal and Ahringer libraries. RNAi bacteria were grown to log phase, then seeded onto MYOB plates containing 1 mM IPTG and 25 ug/ml carbenicillin, and incubated at room temperature for 24 hours. To silence the target genes, 10-15 mid-L4 hermaphrodites were picked and grown on RNAi plates at 25°C for 24-36 hours before imaging.

### Live imaging of animal gonad and cortical granule exocytosis

For imaging the gonad, animals were immobilized on 7% agar pads with an anesthetic (0.1% tricaine and 0.01% tetramisole in M9 buffer or 0.01% levamisole in M9 buffer). DIC images wacquiere captured with a Nikon 60X 1.2 NA water objective and a 0.5 μm z-step size; 15-20 z-planes were acquired. For imaging meiosis I embryos, gravid animals were dissected, and the embryos were mounted in blastomere culture media using a hanging drop chamber ^56,57^.

### CRISPR design

The repair template design followed the standard protocols ^58,59^. Approximately 20 young gravid animals were injected with the prepared CRISPR/Cas9 injection mix as described in the literature ^59^. *pezo-1 NΔ* and *pezo-1 CΔ* mutants were generated by CRISPR/Cas9 mixes that contained two guide RNAs at the flanking regions of the *pezo-1* coding regions. Heterozygous *pezo-1* deletion animals were first screened by PCR and then homozygosed in subsequent generations. All homozygous animals edited by CRISPR/Cas9 were confirmed by Sanger sequencing. The detailed sequence information for the repair template and guide RNAs is listed in the supplemental information and in previous studies ^25,44^.

### Microscopy

Live imaging was performed on a spinning-disk confocal system using a Nikon 60X 1.2 NA water objective, a Hamamatsu C15440 ORCA-Fusion BT Digital camera, and a Yokogawa CSU-X1 confocal scanner unit. Nikon’s NIS imaging software was applied to capture the images. Images were acquired and analyzed by Nikon’s NIS imaging software and ImageJ/FIJI Bio-formats plugin (National Institutes of Health) ^60,61^.

### The microinjection of fluorescein-labeled MSP into *fem-1(hc17)* hermaphrodites

The microinjection of 101.6 μM NHS-Fluorescein-labeled MSP-142 into adult *fem-1(hc17)* hermaphrodites (Day 1-2, 24-36 hours post mid-L4, at non-permissive temperature of 25°C) was performed as previously described ^62^. The injected worms recovered for 4 hours on MYOB plates with OP50 food before imaging. The acquisition of GFP and DIC images was performed using our confocal imaging system (see above) with a 0.5 μm z-step size and 15-20 z-planes.

### Statistics

Statistical significance was determined by p-value from an unpaired 2-tailed t-test. P-values: **** = <0.0001. Both the Shapiro-Wilk and Kolmogorov-Smirnov normality tests indicated that all data follow normal distributions. P-values: n.s. = not significant; * = <0.05; ** = <0.01, *** = <0.001; **** = <0.0001.

## Data Availability

All data necessary to confirm the findings of this study will be made fully available upon publication, including in the manuscript, supplementary files, and public repositories. All *Caenorhabditis elegans* strains generated in this study will be deposited and made publicly available through the Caenorhabditis Genetics Center (CGC).

## Interest of Conflicts

The authors have no conflicts of interest to declare.

## Funding

The project was supported by an NIH Pathway to Independence Award (K99/R00), 1K99 GM145224-01(X.F.B.). National Institute of General Medical Sciences/National Institutes of Health (R00GM145224 and R35GM162564-01 to X.F.B.), and the UF Startup fund for the Bai lab.

## Acknowledgments

We thank the *Caenorhabditis* Genetics Center, which is funded by the National Institutes of Health Office of Research Infrastructure Programs (P40OD010440), for providing strains for this study. We especially thank the UF worm clubs and Drs. Erin Cram, Harold Smith, Chenshu Liu, and Shaohe Wang for their critical input on the project and feedback on the manuscripts. We thank Dr. David Greenstein’s generous gift of the MSP reagents. Lastly, we thank all members of the Bai lab for providing feedback and suggestions on our investigations, as well as for helping to prepare lab reagents and conduct basic data analysis.

---

**Supplemental Figure 1. Germline-specific degradation of PEZO-1 affects vesicle trafficking.**

(A) Schematic diagram showing the working mechanism of the tissue-specific Auxin-Inducible Degradation (AID) system. The target protein (PEZO-1::AID) undergoes targeted polyubiquitination and subsequent proteasomal degradation only in cells co-expressing the F-box substrate recognition protein TIR-1 driven by a tissue-specific promoter (*pie-1p* for germline; *eft-3p* for pan-soma). (B–C) Representative images of adult hermaphrodite gonads expressing GFP::PEZO-1::Degron and germline-specific Ppie-1::tir-1::mRuby. Basal expression in the control animal (B) shows vesicular accumulation in the proximal oocytes (labeled -3, -2, -1), whereas exposure to 2 mM auxin (IAA) (C) selectively abolishes the germline signal while preserving somatic expression of GFP::PEZO-1::AID. (D) Quantitative analysis of the relative GFP fluorescence ratio in the germline-specific AID strain with both control and auxin treatment. (E–F) Representative images of animals expressing GFP::PEZO-1::Degron and somatic Peft-3::tir-1::mRuby. The control animal (E) maintains vesicular populations in the germline in both control and auxin treatment. However, upon treatment with 2 mM auxin (F), the somatic channel fractions (including intense signals within the spermatheca) are significantly reduced. Scale bars are indicated in each panel. P-values: ****: p <0.0001 (t-test). Scale bars are indicated in each panel.

**Supplemental Figure 2. PEZO-1 does not traffic through early endosomes but dynamically co-localizes with the recycling receptor VAB-1.**

(A–B) Representative images of the strain co-expressing GFP::PEZO-1 (green) and the early endosomal marker mCherry::RAB-5 (magenta) across the distal (A) and proximal (B) hermaphrodite germline cortex. Yellow arrows in the magnified right inserts highlight completely distinct, non-overlapping vesicular paths. (C–D) While the recycling Ephrin receptor VAB-1::GFP (green) and mScarlet::PEZO-1 (magenta) are overly co-localized at the characteristic ring structures at the cortical regions of the distal (C) and proximal (D) gonad. Yellow arrows in the magnified right insets track individual dual-positive transport vesicles. Scale bars are indicated in each panel.

**Supplemental Figure 3. PEZO-1 and RAB-11 detached following embryonic exocytosis.**

(A–D) Live-imaging of a meiotic embryo co-expressing mScarlet::PEZO-1 (magenta) and the trafficking regulator RAB-11::GFP (green). Prior to the exocytic wave, RAB-11::GFP and mScarlet::PEZO-1 co-occupy the cortical granules (yellow arrows, A–C). Immediately following vesicle fusion at anaphase I onset (03:33), the RAB-11::GFP delivery signal rapidly dissociates from the membrane and clears into the cytoplasm, whereas the mScarlet::PEZO-1 channel remains stably integrated into the remodeled embryonic plasma membrane (red arrows, D). Timestamp indicates minutes: seconds.

**Supplemental Figure 4. Pathogenic mutations impair PEZO-1 expression at the pharyngeal-intestinal valve.**

(A–G) Representative images show the pharyngeal-intestinal valve tissue in adult animals expressing endogenously tagged wild-type GFP::PEZO-1 (A) and in animals carrying pathogenic mutations in the *C. elegans* pharynx. Panels show the pore helix mutant R2373C (B), R2373H (C), the C-terminal domain (CTD) mutant R2405L (D), R2405P (E), and the corresponding double-mutant combinations R2373C; R2405L (F), and R2373H; R2405Q (G). Red arrows mark pharyngeal structures, and yellow arrows indicate the pharyngeal-intestinal valve. (H) Quantification of the relative pharyngeal-intestinal valve GFP::PEZO-1 signal intensity across wild-type and all missense mutants. P-values: *: p <0.05; **: p <0.01; ***: p <0.001; ****: p <0.0001 (One-Way ANOVA with Dunnett’s post-hoc test). Scale bars are indicated in each panel.

**Supplemental Figure 5. Pathogenic mutations disrupt the stable accumulation of PEZO-1 channels in the spermatheca.**

(A–G) Representative confocal images of adult hermaphrodite proximal gonads focused on the spermathecal cells (highlighted by yellow dashed boxes) in wild-type GFP::PEZO-1 (A), pore helix variants R2373C (B) and R2373H (C), CTD variants R2405L (D) and R2405P (E), and the double-mutant combinations (F–G). Missense mutations within the C-terminal structural domains cause a profound loss of local structural stability, preventing channel accumulation at the spermathecal cells. (H) Quantification of the spermathecal GFP::PEZO-1 fluorescence intensity ratio across the indicated genotypes. P-values: *: p <0.05; **: p <0.01; ***: p <0.001; ****: p <0.0001 (One-Way ANOVA with Dunnett’s post-hoc test). Scale bars are indicated in each panel.

**Supplemental Figure 6. Genetic depletion of the genetic suppressor allele *gex-3* does not compromise reduced germline PEZO-1 expression in *pezo-1*[R2405P] mutant.**

(A–D) Representative images of confocal micrographs of adult proximal gonads evaluating the structural reliance of PEZO-1 trafficking on the SCAR/WAVE complex component GEX-3. Strains expressing wild-type GFP::PEZO-1 (A–B) or the destabilized disease variant GFP::PEZO-1(R2405P) (C–D) were treated with either control RNAi (A, C) or *gex-3*(RNAi) (B, D). Yellow dashed boxes outline the spermatheca. (E–F) Quantitative analysis of the relative GFP::PEZO-1 fluorescence intensity ratio within the oocyte cytosol (E) and the somatic spermatheca tissue (F) under control versus *gex-3*(RNAi) conditions. RNAi knockdown of *gex-3* significantly reduced channel accumulation at the somatic spermatheca. P-values: *: p <0.05 (One-Way ANOVA with Dunnett’s post-hoc test). Scale bars are indicated in each panel.

## The list of strains used in this study

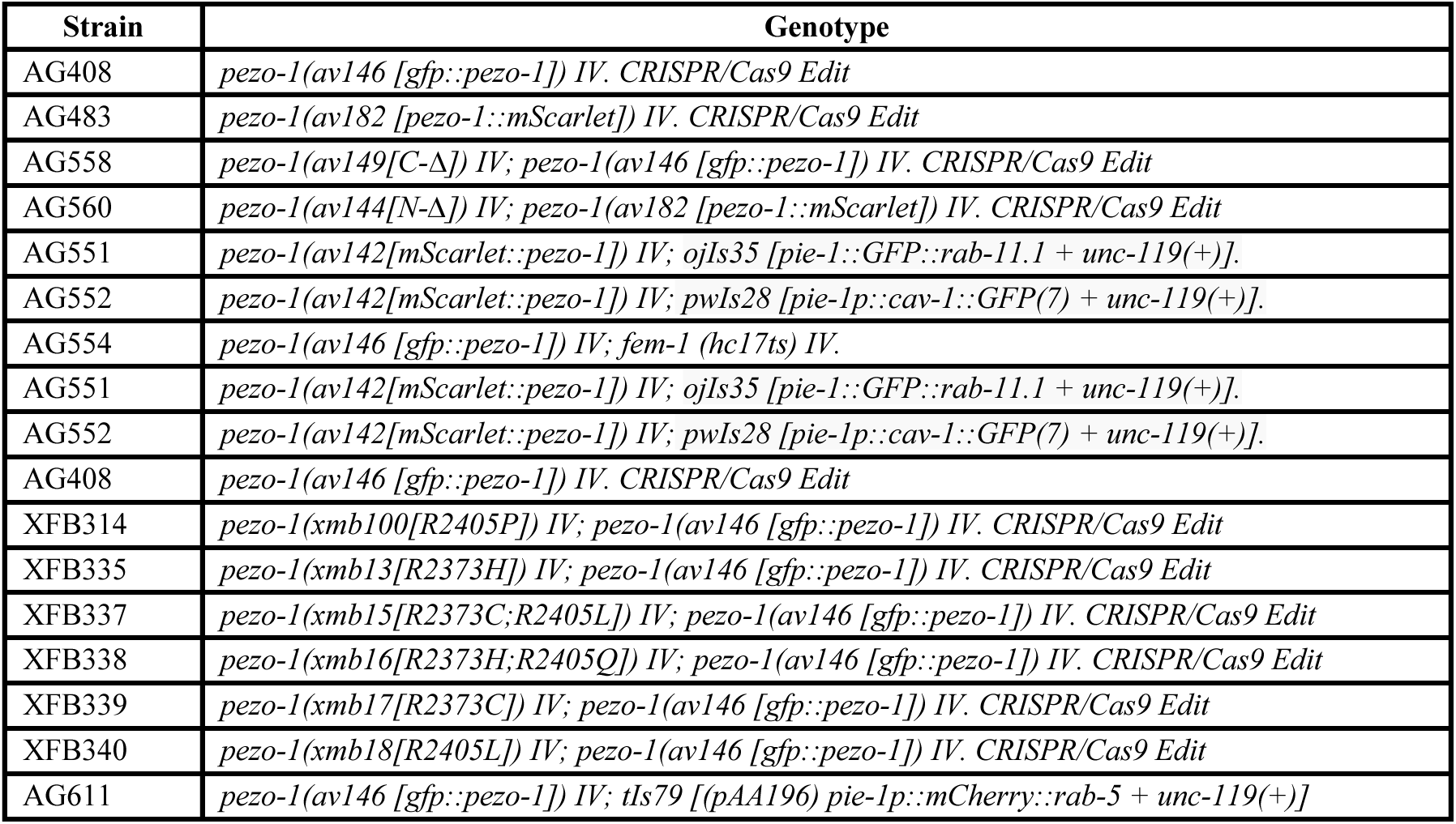

**CRISPR Design and Sequence Information**

*pezo-1C-*Δ CRISPR/Cas9 editing design:

N-terminus Guide RNA: 5’ CGGTGGCAGCGTACATTATC 3’

C-terminus Guide RNA: 5’ CACCAGCGACACTCATCGAA 3’

Genotype Forward Primer: 5’ AATCTGACTTGTGCCCTCCG 3’

Genotype Reverse Primer: 5’ AATCAGGCGAGCAGTGAGAG 3’

Genotype Internal Primer 5’ TCCACAGTCAATTCCTGCGT 3’

Repair template: 5’ tccagtctcccatatttattttttttctgttccagTAGATAAGTAAGAGCAAAAAGAAGCAAGAATAA 3’

*pezo-1N-*Δ CRISPR/Cas9 editing design:

N-terminus Guide RNA: 5’ ACACAGCAACAACAGAATGA 3’

C-terminus Guide RNA: 5’ TGGGGGTGTTGCAGTGGCTA 3’

Genotype Forward Primer: 5’ GCGGTAAATCTGAATCGGTGG 3’

Genotype Reverse Primer: 5’ TTGGAAAAGCAGGCACAACC 3’

Genotype Internal Primer 5’ CGATCCAGCGTGGATGAACT 3’

Repair template: 5’ atctgaatcggtggtcgtaacacagcaacaacaga**g**tttgacacattttccgttgagacttgaaaaatag 3’

*pezo-1*[R2373C] CRISPR/Cas9 editing design:

Guide RNA: 5’ AAGATTCCACGAACCAGACC 3’

Repair template: 5’ TCTGTGAACATGACAGTGCTTGGAGATGTGGTGAAAATACCACACACGAGACctggaagaaaatgagactggtaagaatttcctaa 3’

Restriction Enzyme: BsaI

*pezo-1*[R2373H] CRISPR/Cas9 editing design:

Guide RNA: 5’ AAGATTCCACGAACCAGACC 3’

Repair template: 5’ TCTGTGAACATGACAGTGCTTGGAGATGTGGTGAAAATACCATGGACAAGACctggaagaaaatgagactggtaagaatttcctaa 3’

Restriction Enzyme: NcoI

*pezo-1*[R2405L] CRISPR/Cas9 editing design:

Guide RNA: 5’ CTATTTGGTTCGAGAAGCGA

Repair template: 5’ GATCATCTTCTCAAAATTTGTCTCGACATCTATTTAGTACTAGAAGCGAAAGACTTCATGTTGGAGCAGgtaattatttagtttta 3’

Restriction Enzyme: ScaI

*pezo-1*[R2405P] CRISPR/Cas9 editing design:

Guide RNA: 5’ CTATTTGGTTCGAGAAGCGA 3’

Repair template: 5’ CATCTTCTCAAAATTTGTCTCGACATCTATTTGGTACCAGAAGCGAAAGACTTCATGTTGGAGCAGgtaattatttagtttta 3’

Restriction Enzyme: KpnI

*pezo-1*[R2405Q] CRISPR/Cas9 editing design:

Guide RNA: 5’ CTATTTGGTTCGAGAAGCGA 3’

Repair template: 5’ CATCTTCTCAAAATTTGTCTCGACATCTATTTGGTGCAGGAAGCGAAAGACTTCATGTTGGAGCAGgtaattatttagtttta 3’

Restriction Enzyme: BsgI

## Notes

### Competing Interest Statement

The authors have declared no competing interest.

