## Supplemental Figure 1-6 for "Regulation of Mechanosensor PIEZO Channel Trafficking in *C. elegans* Germline"

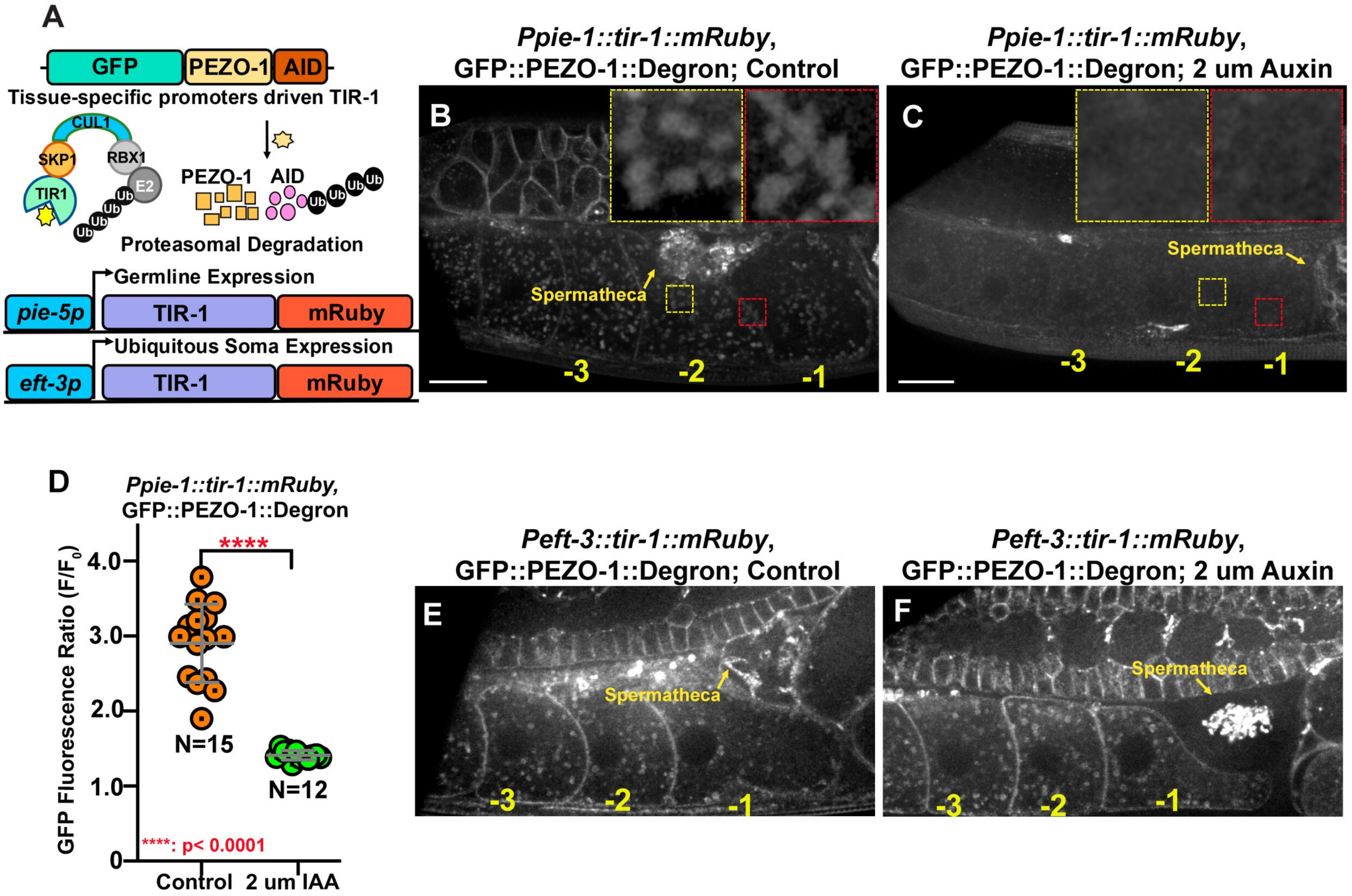

### Supplemental Figure 2

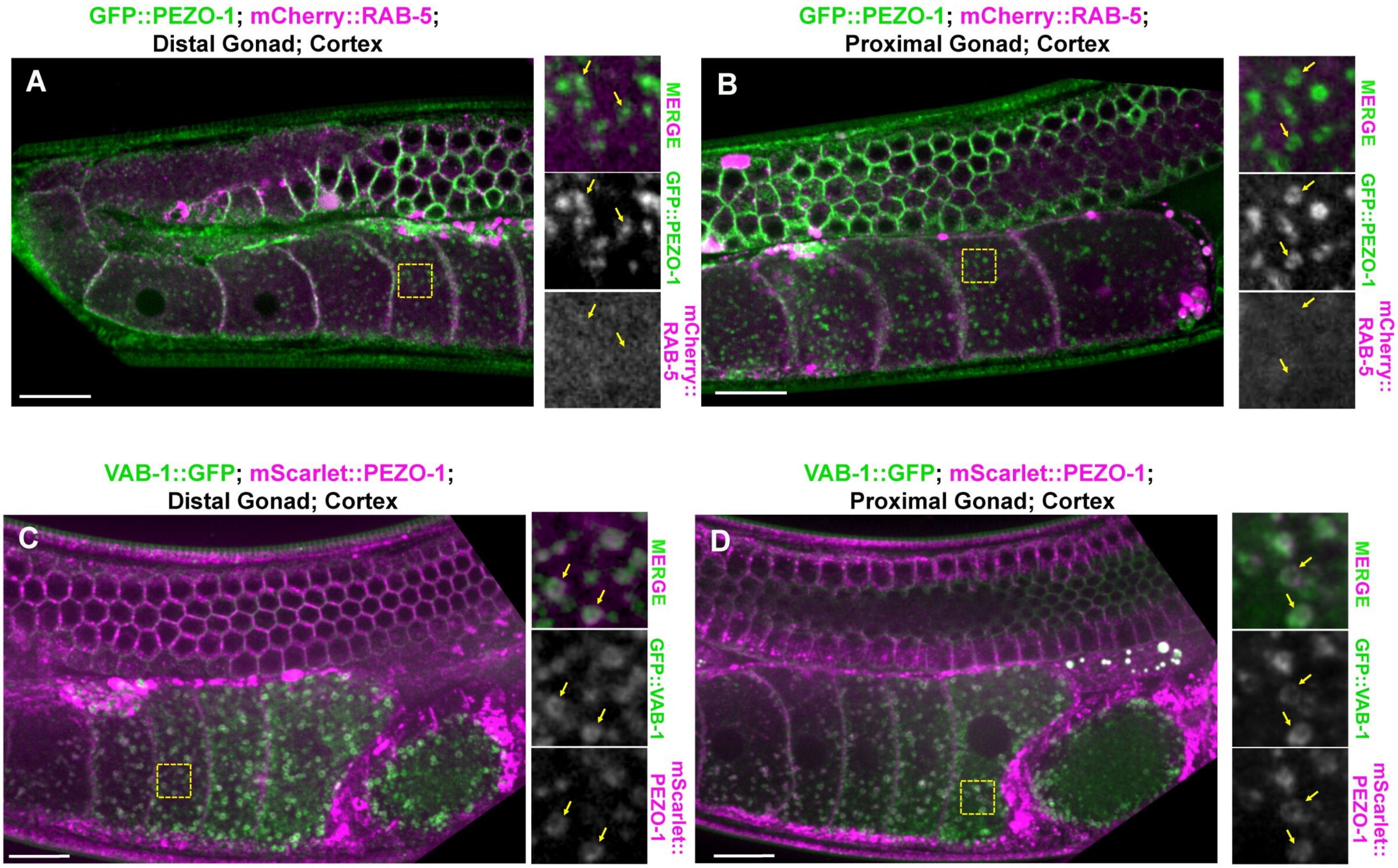

#### Supplemental Figure 3

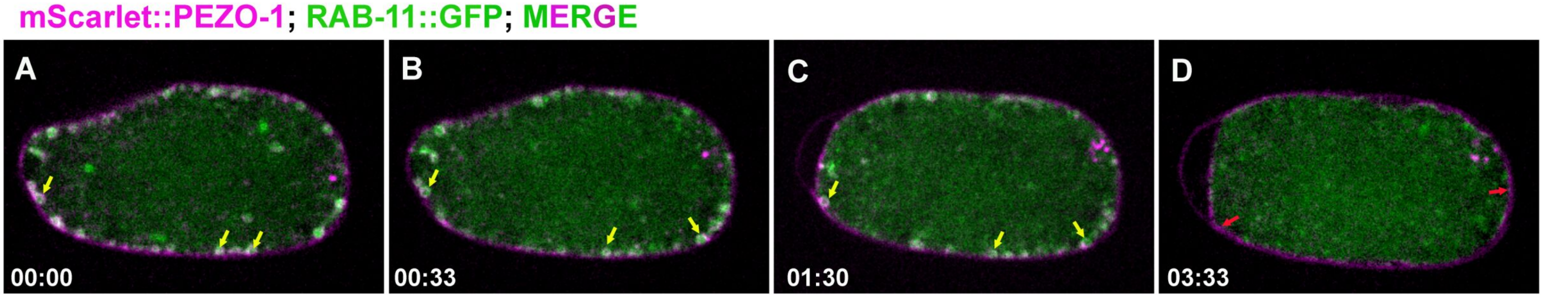

#### Supplemental Figure 4

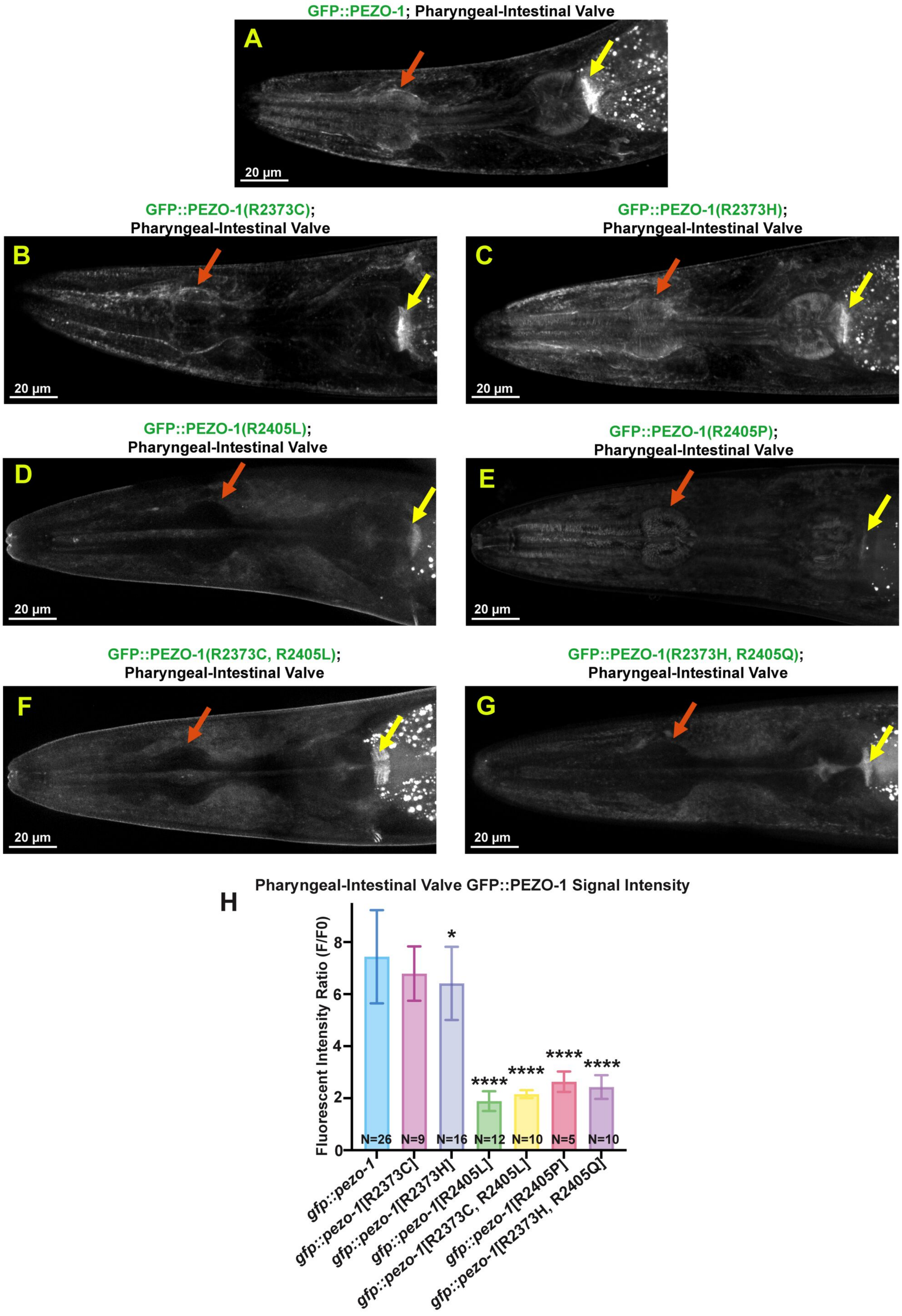

Supplemental Figure 5

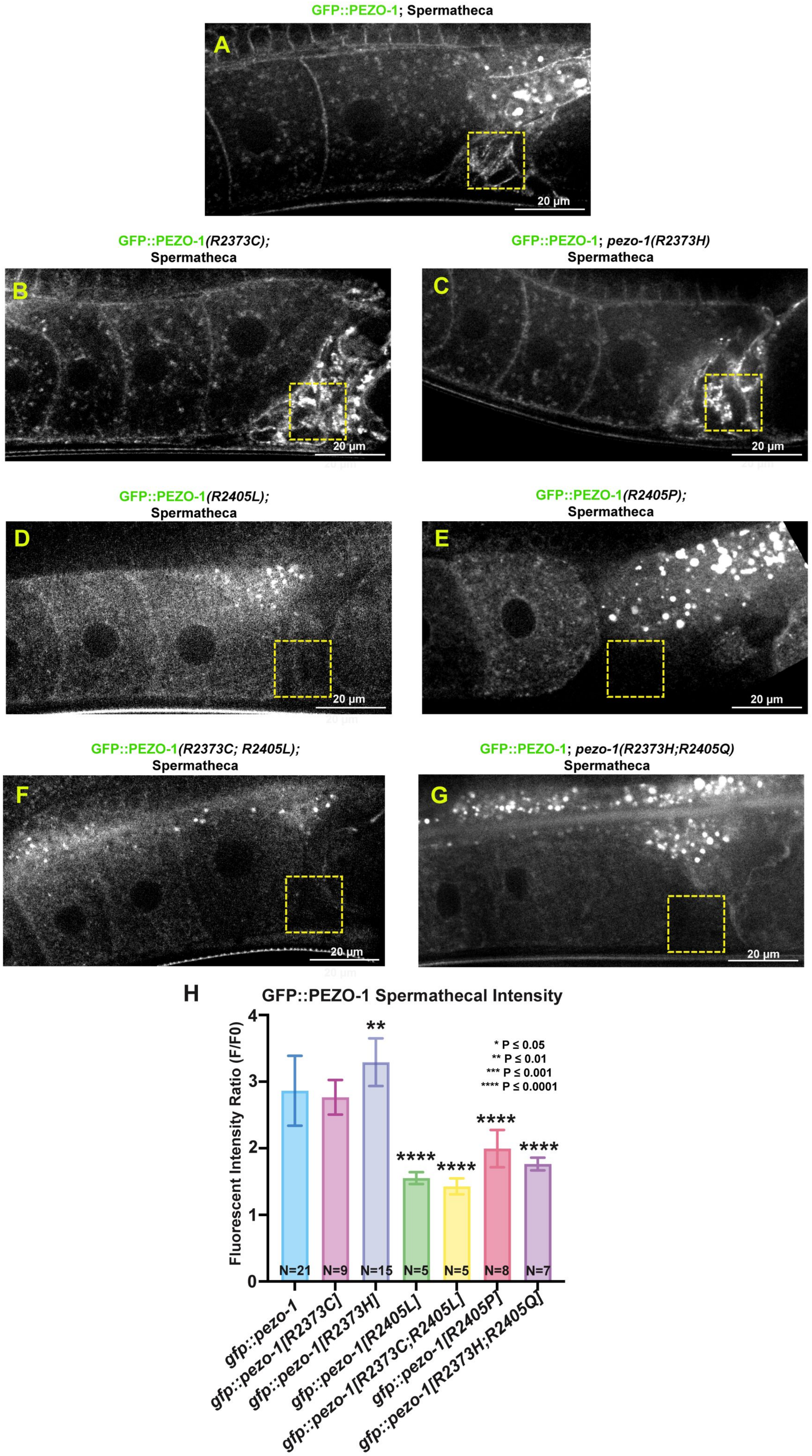

#### Supplemental Figure 6

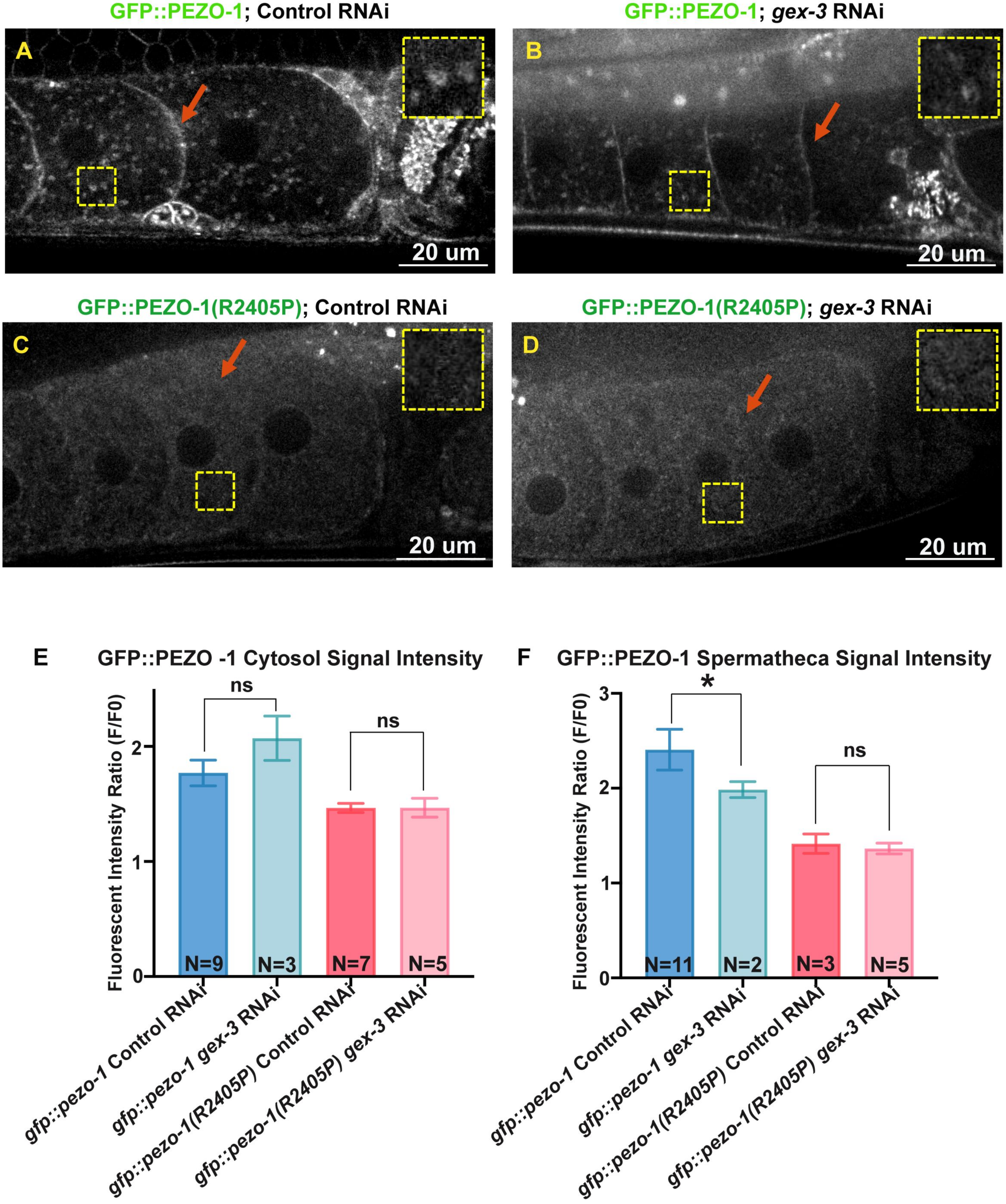
